## Supplementary material for "Sulfide-responsive transcription control in *Escherichia coli*": Supp_Figs

### Volcano-plot

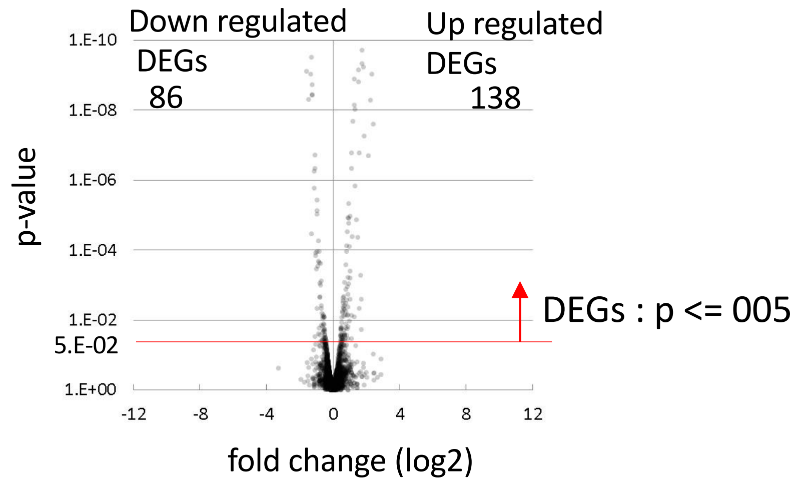


AC : Aerobic condition
SA : Semi-aerobic condition
SA+$Na_2S$ : Semi-aerobic condition + Na2S

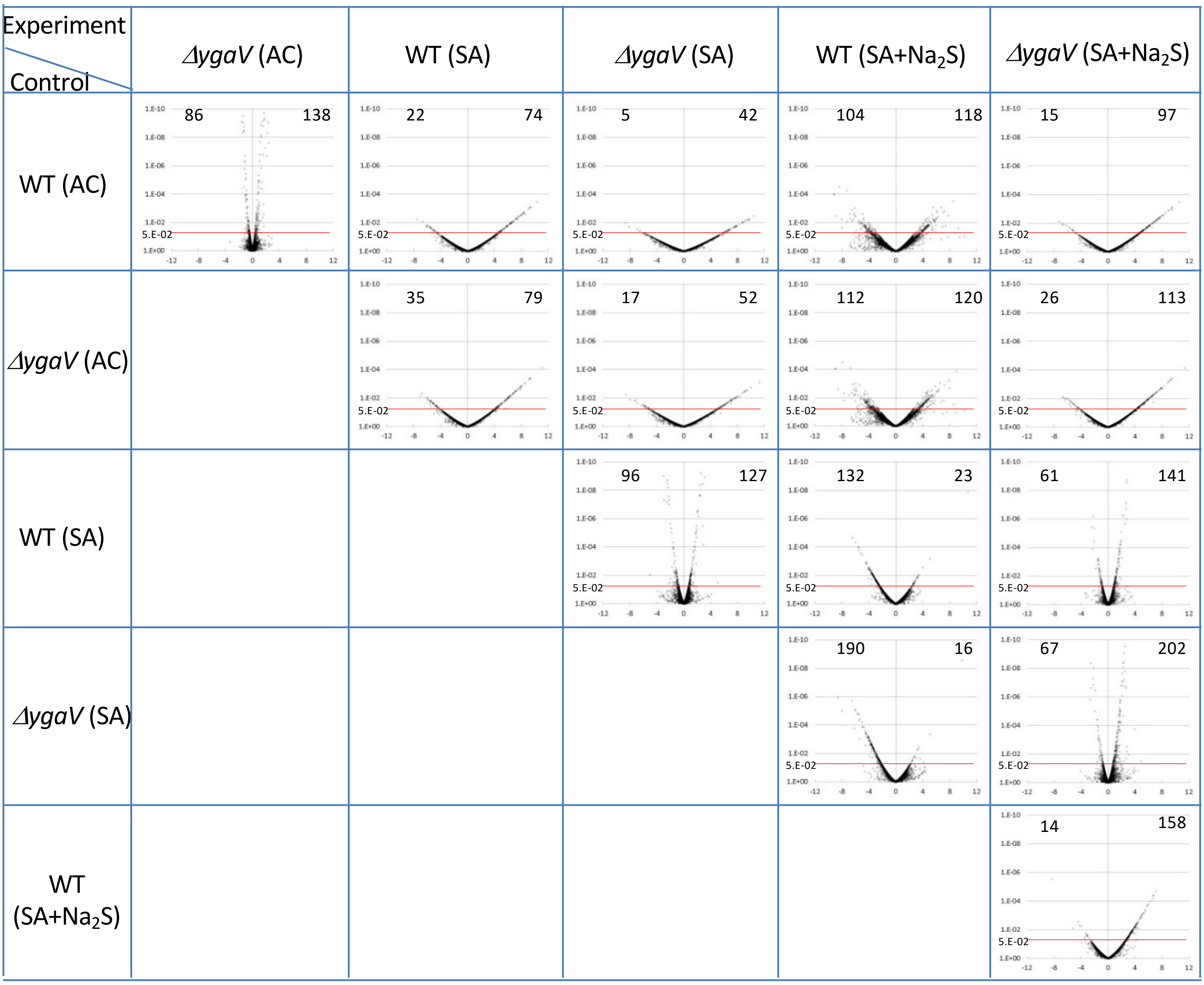


**Supplemental Fig. S1** Volcano plots for upregulated and downregulated transcripts in WT and *ΔygaV* mutant grown under aerobic (AC) and semi-aerobic (SA) conditions with (+$Na_2S$) or without 0.2 mM $Na_2S$.

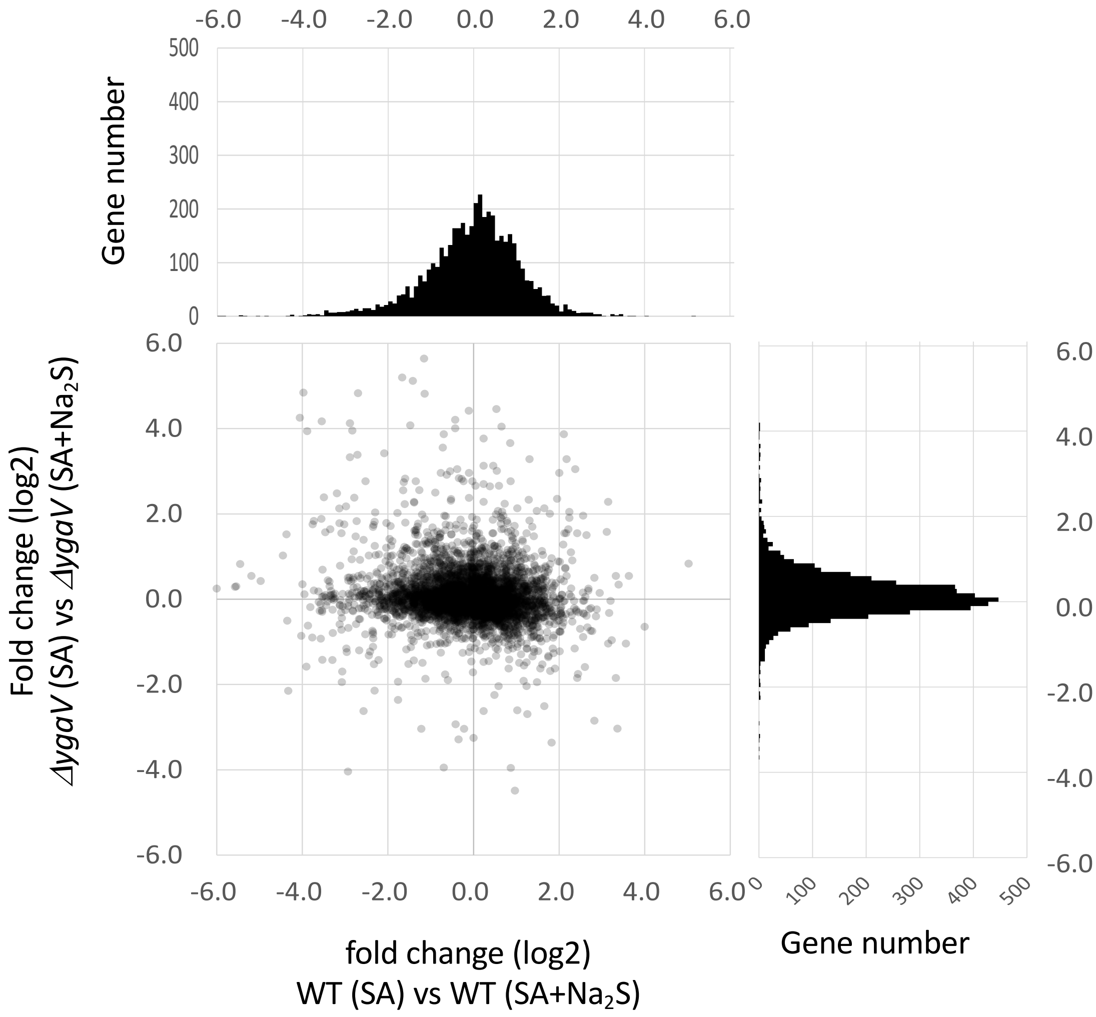


**Supplemental Fig. S2** Scatter plot depicting distributions of the expression changes between WT (SA) and WT (SA+$Na_2S$) (x-axis, top marginal histogram) against the expression changes between *ΔygaV* (SA) and *ΔygaV* (SA+$Na_2S$) (y-axis, right marginal histogram).

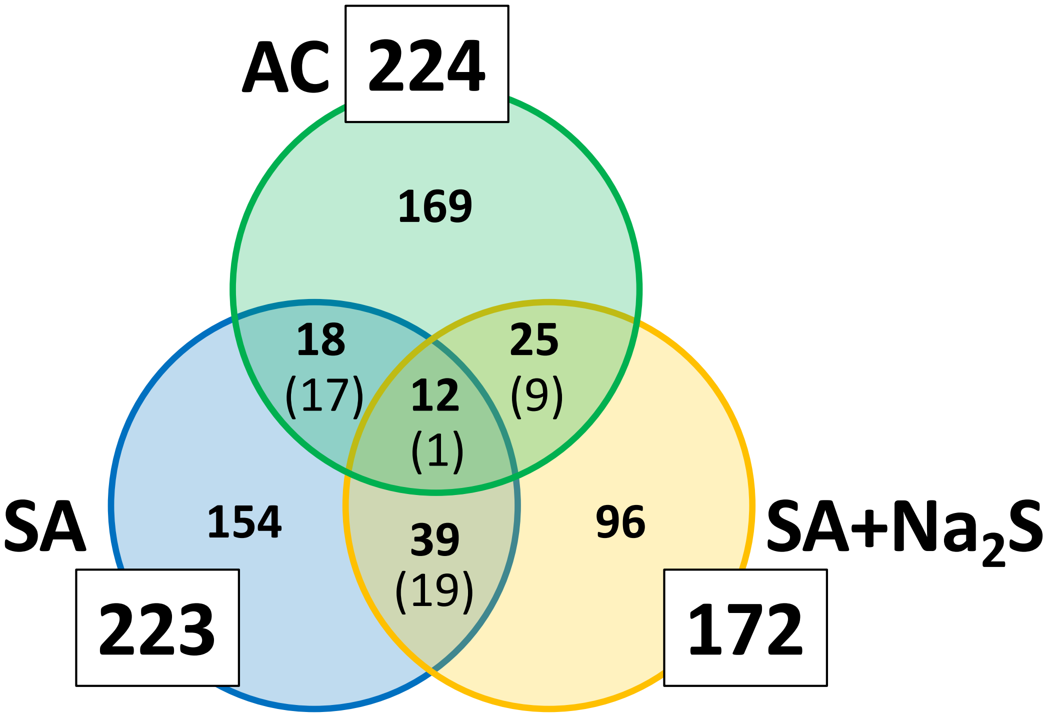


**Supplemental Fig. S3** Venn diagrams depict the overlap between genes differentially expressed in the *ΔygaV* mutant compared to WT, grown under aerobic (AC) and semi-aerobic (SA) conditions, with (+$Na_2S$) or without 0.2 mM $Na_2S$. The numbers in brackets indicate the number of genes with concordant increased or decreased expression in the two comparisons.
